# openNAU: An open-source platform for normalizing, analyzing, and visualizing untargeted metabolomics data

**DOI:** 10.1101/2022.08.31.506116

**Authors:** Qingrong Sun, Qingqing Xu, Majie Wang, Yongcheng Wang, Dandan Zhang, Maode Lai

## Abstract

**Motivation:** As an important part of metabolomics analysis, untargeted metabolomics has become a powerful tool in the study of tumor mechanisms and the discovery of metabolic markers with high-throughput spectrometric data which also brings great challenges to data analysis from the extraction of raw data to the identification of differential metabolites. To date, a large number of analytical tools and processes have been developed and constructed to serve untargeted metabolomics research. The different selection of analytical tools and parameter settings lead to varied results of untargeted metabolomics data. Our goal is to establish an easily operated platform and obtain a repeatable analysis result.

**Results:** We used the R language basic environment to construct the preprocessing system of the original data and the LAMP (Linux + Apache + MySQL + PHP) architecture to build a cloud mass spectrum data analysis system. An open-source analysis software for untargeted metabolomics data (openNAU) was constructed. It includes the extraction of raw mass data and quality control for the identification of differential metabolic ion peaks. A reference metabolomics database based on public databases was also constructed. Finally, a complete analysis system platform for untargeted metabolomics was established. This platform provides a complete template interface for the addition and updating of the analysis process, so we can finish complex analyses of untargeted metabolomics with simple human-computer interactions.

**Availability and Implementation:** The source code can be downloaded from https://github.com/zjuRong/openNAU.

## Introduction

Untargeted analysis is a research method that performs a high-throughput quantitative analysis of unknown metabolites by comparing the metabolites of a control group and an experimental group to find differences in their metabolic profiles(Schrimpe-Rutledge, et al., 2016). In the research process, untargeted metabolomics can be used to screen metabolites with differences in the disease development process, and then further qualitative and quantitative verification through targeted metabolomics can be used for early diagnosis and prognosis assessment. The main analytical techniques used for data collection are nuclear magnetic resonance (NMR), mass spectrometry (MS), chromatography, and spectroscopy. The complementarity of analytical techniques solves the problem of the difference in the scope, composition, and nature of the samples tested(Dai, et al., 2010; Gao, et al., 2008). NMR, LC- and GC-MS are the three most commonly used data acquisition techniques and have become the mainstream of metabolomics research.

The preprocessing and postanalysis of untargeted metabolomics data play an important role in metabolomics research. In the preprocessing of metabolomics data, Isotopologue Parameter Optimization (IPO) (Libiseller, et al., 2015), XCMS(Smith, et al., 2006), CAMERA(Kuhl, et al., 2010), Workflow4Metabolomics(Giacomoni, et al.), MetaboAnalyst(Chong, et al., 2018), and MetFlow(Thin, et al., 2020) are need as quantification tools of ion peaks. The main function of IPO is to optimize parameters in XCMS so that XCMS or CAMERA can retain valuable ion peak information as much as possible under optimized parameter settings. XCMS and CAMERA quantify the ion peaks of the raw data based on algorithms, but CAMERA is more valuable for labeling the composition of ions. The quantization matrix of ion peak intensity required by a study can be obtained through the above processing, and the missing values in the quantized ion peak can be filled based on MICE(Buuren and Groothuis-Oudshoorn, 2010). After the preprocessed data are obtained, Normalyzer(Chawade, et al., 2014), Norm ISWSVR(Ding, et al., 2022) and MSPrep(Grant, et al.) can be used for data correction and normalization, and finally, statistical analysis and model construction of normalized ion peak intensity differences can be carried out.

Surely, the optimization of parameters and quantification of ion peaks in the raw data extraction process have been recognized. There are many methods to fill the quality control of ion intensity, but there are some differences. After the raw data pretreatment process, the correction of ion intensity also means that although many methods can be used to normalize data, there is no unified standard, so the normalized data cannot obtain consistent analysis results. In addition, the data batch effect and instrument error are objective factors that affect the conclusions. We aim to strengthen the normalization of data quality control, build a complete analysis process, and simplify the complex parameter setting and tool selection in the analysis process.

## Methods

### 1. Overall design

We built the whole analysis process based on the R programming language and LAMP (Linux + Apache + MySQL + PHP) architecture. The software consists of two subsystems: preprocessing of raw data (MetaQC) and metabolomics analysis and resources central (MARC) (Figure 1). MetaQC uses the Shiny(Chang, et al., 2017) module in the R language to realize the visualization operation of the analysis process, and inserts the extraction and analysis program of raw data into the visualization operation process. MARC is based on a cloud server for data analysis and processing. This section is divided into the front end, back end, and cloud analysis strategy. The front end is based on the bootstrap framework (www.bootcss.com) for the front-end layout and communication with the back-end. The back-end uses PHP for database access, file operation, and data transmission with the front-end. The cloud analysis strategy uses a socket as the interface to realize the call of cloud analysis software, the acquisition of tasks and the corresponding data analysis, and the analysis results through the intermediate log file for mutual result authentication and process identification.

**Figure 1.**
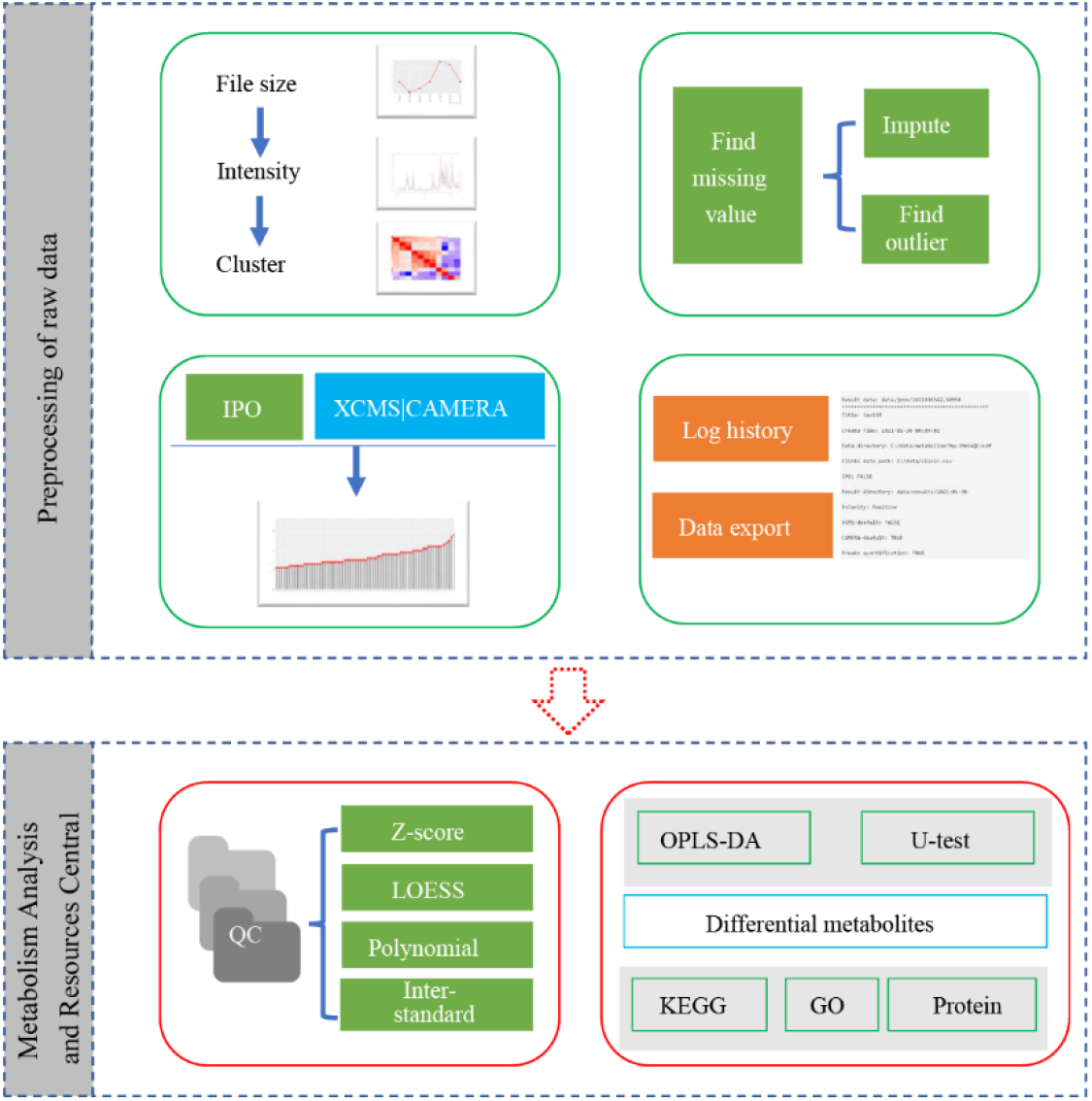
Workflow of the software.

### 2. Function design

We recommend using Agilent’s Masshunter Qualitative Analysis and DA Reprocessor software to create quantitative parameter files for raw data and convert bulk data. In the parameter file, the main parameter that needs to be customized is the ion intensity threshold setting. Here, we set it to 500-1000. In this way, some noise ion peaks can be screened out. Finally, the raw data are transformed into a readable data structure, and the size of exported files is compared to ensure that the file size is approximately 10 MB.

#### 2.1 Parameter optimization and ion peak quantization

We used IPO software to optimize the parameters of the screened data and then combined it with CAMERA or XCMS software for analysis after obtaining the optimized parameters. Of course, users can also directly use XCMS and CAMERA for quantization based on the parameters set in advance. In our software, the default parameters are FWHM =10, method = “obiwarp” and plotType = “deviation” when XCMS and CAMERA are directly run to obtain ion peaks. Optimization parameters during IPO operation will refine the default parameters and optimize the ion peak extraction method, retention time correction method, and ion peak grouping method(Libiseller, et al., 2015). The selection of the ion peak extraction methods centWave and matchedFilter needs to be determined according to the research data. CentWave was selected for high-resolution data, and matchedFilter was selected for low-resolution data.

#### 2.2 Quality control and missing value filling

A custom R program and mvoutlier(Filzmoser P, 2018) package are used to control the quality of the samples and ion peaks. First, the missing data of ion peaks and samples are removed based on the custom R program. The threshold in our program is the elimination of samples based on more than 50% ionic peaks with 0 intensity and the elimination of ionic peaks by more than 20% samples with 0 intensity. Then, multivariate outlier detection(Filzmoser, et al., 2008) is performed on the above results using mvoutlier software, and the abnormal samples are removed again. The missing values are filled by one of 6 alternative methods (Van Buuren, 2018) based on the MICE package.

#### 2.3 Data normalization

This function conducts intrabatch and interbatch correction of quantified metabolic ionic peak data, which is a benefit to later data analysis. First, the data are calibrated regarding internal standard ions so that baseline differences between samples can be eliminated. The data are then corrected interbatches based on the quality control samples. In our software, MetaboQC (Calderón-Santiago, et al., 2017), SVA (Jeffrey T. Leek, 2020), and a custom R program was used to normalize the quantized ion peak intensity data.

First, the data are normalized by using formulas (1), (2), and (3) to eliminate the intragroup differences and intergroup differences and reduce the influence of outlier values (showed in Formula 1). In the formulas, I(peak_ij_) refers to the peak intensity of the i-th ion peak of the j-th sample; I(peak_j_) refers to the ion peak intensity value of the internal standard of the j-th sample; Mean(Q_i_) refers to the mean intensity of the i-th ion peak in all quality control samples; σ(Q_i_) refers to the standard deviation of the i-th ion in all quality control samples; Min(peak_i_) and Max(peak_i_) refer to the minimum and maximum value of the i-th ion peak in all experimental samples, respectively.

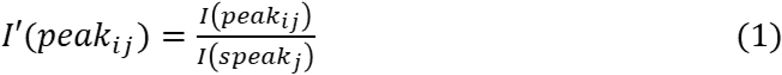

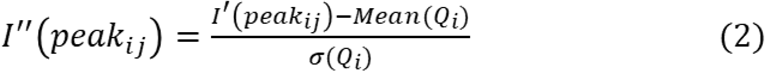

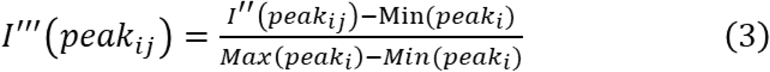

In addition, we used the polynomial correction function of the MetaboQC software to correct the data based on the trend of quality control samples. At the same time, the function of batch correction of expressed data concerning the SVA tool can be applied to eliminate the batch effect of metabolic ion strength values.

#### 2.4 Differential analysis

The difference analysis of metabolic ions requires a customized program and the ropls software package for comprehensive analysis. Then the results are verified by orthogonal partial least square discriminant analysis (OPLS-DA) (Boccard and Rutledge, 2013) and a custom program to obtain the final differential metabolic ionic peaks. The normalized ion peak data are used for a Mann-Whitney U test or T-test analysis, and the *p* value or adjusted *p* value (FDR) is outputted. The default thresholds are FDR<0.05 or P<0.05, Fold change > 2 or <0.5, and VIP (Variable Importance in the Projection) >1.

#### 2.5 Metabolites identification

First, we constructed an Integration of reference metabolomics (refMet)(Fahy and Subramaniam, 2020) database based on The human metabolome database(HMDB) (Wishart, et al., 2007), PubChem(Yanli, et al., 2009), The Small Molecule Pathway Database (SMPDB)(Alex, et al., 2010), Metabolomics Workbench(Sud, et al.), EzCatDB(Nagano, 2005), Reactome(David, et al., 2014), AllCCS(Zhou, et al., 2020), PathBank(Ho, et al., 2006), MassBank(2010), NPAtlas(van Santen, et al., 2019), and MSigDB(Liberzon, et al., 2011). Then we matched the different metabolic ions based on the refMet database. The charge of ions needs to be removed from the nucleo-mass ratio (m/z), and then the median m/z of ions needs to be set as the central value. Finally, the qualified metabolic ions in refMet were screened with ±10 ppm as the error range. Of course, this range can be set as needed.

#### 2.6 Pathway/GO enrichment analysis

In this section, we constructed our metabolic pathway database based on refMet data. Moreover, clusterProfiler (Yu, et al., 2012) was used to conduct pathway enrichment analysis for the obtained differential metabolic ions or their target genes. The GSVA software (Hnzelmann, et al., 2013) can be used to score the pathway for the obtained differential metabolic ions to analyze the correlation between metabolic ions and pathways.

## Results

The untargeted metabonomic analysis platform is divided into two modules: a localized data preprocessing module (Figure 2A) and a cloud server module (Figure 2B). Both modules realize visual distribution operation and visual display of results. Meanwhile, the two modules can be used for data API, increasing the utility of the tool. MARC data normalization of MARC is based on internal standard ions and/or quality control samples and presented based on the framework in Figure 2C. The overall mean values and standard deviation distribution changes of data before- and after-normalization are compared. In this module, we constructed two data normalization pipelines, including data normalization for experiments containing both internal standard ions and quality control samples or only internal standard ions for the experiment. Figure 2D shows the framework for the data analysis process and the statistical approach in each analysis node. In addition, we selected the data in faahKO(Saghatelian, et al., 2004) as an example for the preliminary test for MetaQC and completed the usage document (https://github.com/zjuRong/openNAU/blob/main/Document.pdf). Finally, the data from kidney cancer and colorectal cancer samples in our laboratory were used for further verification.

**Figure 2.**
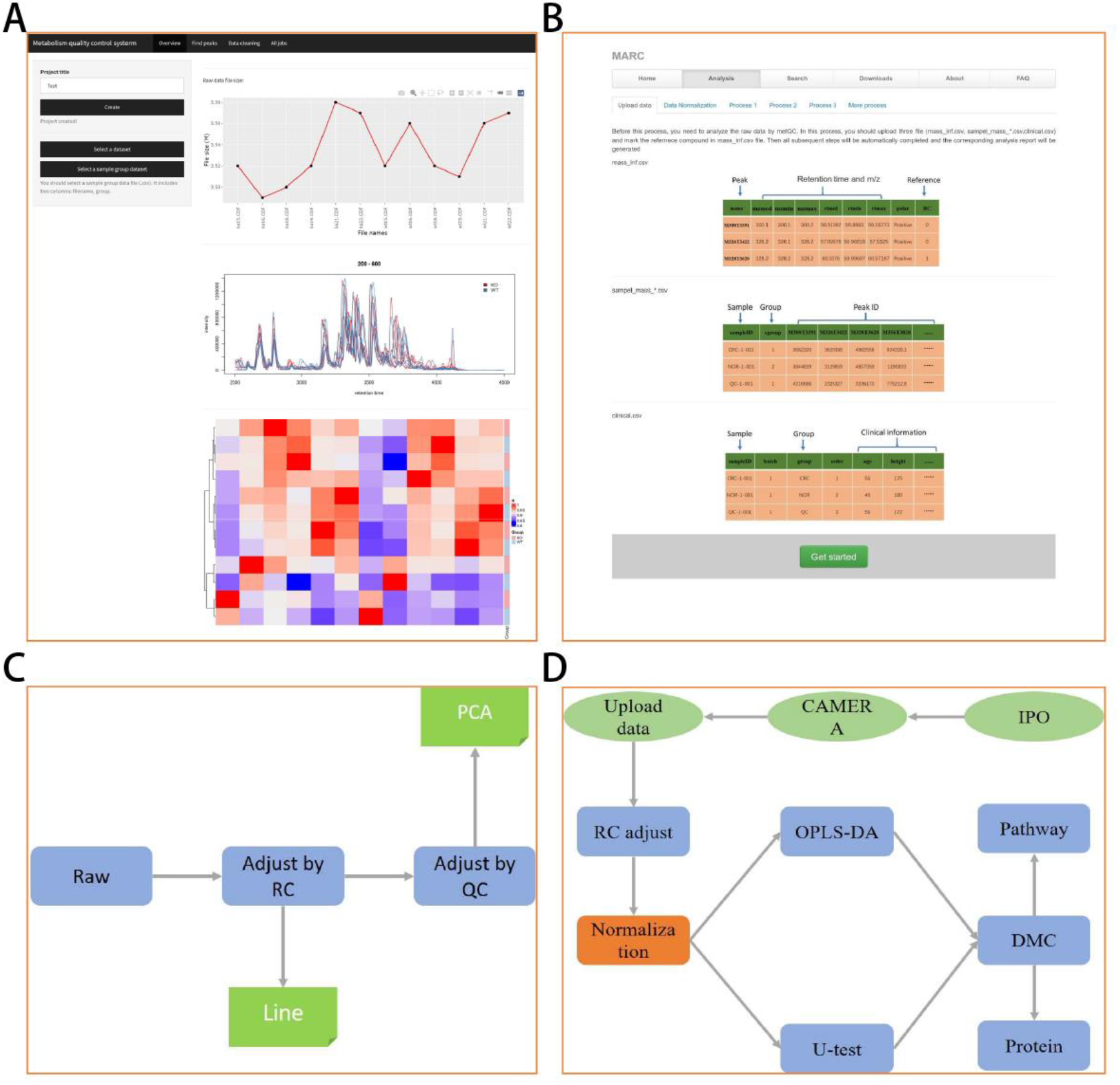
The main page for the analysis platform. A) MetaQC index page. B) MARC index page. C)Frame for data normalization. D) Analysis pipeline for MARC. Raw refers to the uploaded raw peak intensity data, RC refers to the internal standard ion, PCA refers to the principal component analysis, QC refers to the data of the quality control sample, U test refers to Mann-Whitney rank-sum test in the Wilkerson rank-sum test, OPLS-DA refers to orthogonal partial least square discriminant analysis, and DMC refers to different metabolic compounds. In C and D, green color refers to results presentation, blue color refers to the analysis node or methods, reseda refers to the analysis process in MetaQC module and orange-brown refers to all normalization methods (Z score, LOESS, SVA, spoly3 and spoly4).

We collected data from 453 blood samples in our laboratory and some operational errors in these experimental data. In these data, there were 152 healthy control samples, 36 quality control samples, and 265 patients with renal carcinoma (RCC) before (n=154) and after (n=111) surgery. From the MetaQC analysis, we first found the change in peak intensity of all samples with retention time. According to the results, batch 18 samples showed significant enrichment, while other batches of samples showed varying degrees of fluctuation (Figure 3A). In Figure 3B, according to the clustering results of each batch in the heatmap, we can see that multiple samples of a single batch were partly clustered, and the peak intensity of multiple batches of samples was clustered. The above results showed that although there was no difference in peak intensity among batches, there were significant differences among samples from each batch. To find the cause of the fluctuation that interferes with all of the data, we conducted an analysis based on the missing values of data samples and ion peaks and found that only one peak was seriously missing. Then we imputed the missing data and found that there was no difference between before- and after-imputation (Figure 3D). In addition, we detected outliers and found that there were anomalies in many samples. The two most obvious samples were X2241 and X2521 (Figure 3C).

**Figure 3.**
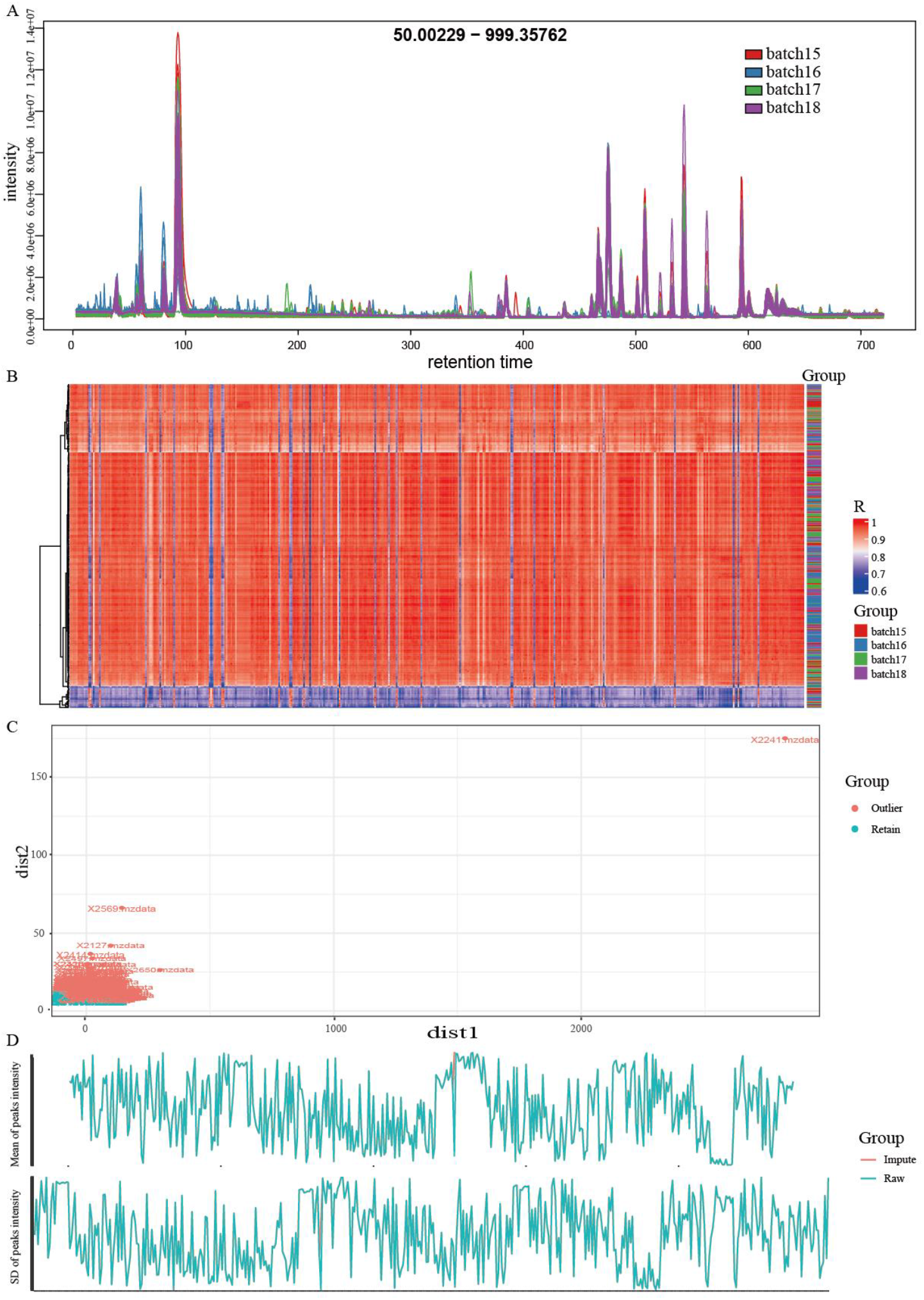
Raw data evaluation and cleaning. A) Line plot for the intensity of peaks with retention time, B) Heatmap for sample distance based on peak intensity, C) Scatter plot for outlier results, D) Mean and SD comparison before- and after-imputation.

According to the above results, we can preliminarily determine that there are some problems in these data. Therefore, we carry out data normalization processing based on MARC, aiming to eliminate the noise of such data. First, we normalized the QC data of the experiment by normalization pipeline 1 in MARC. We found that there were significant differences in QC data among the different groups (Figure 4G and Figure 5D). In addition, we found that raw QC data were affected by the batch effect (Figure 4A) and only the Z-score method corrected the QC data with obvious noise removal (Figure 4C). Among other methods, QC data failed to be corrected by RC (Figure 4B) and Single-Poly3 (Figure 4E) and was overcorrected by LOESS (Figure 4D) and Single-Poly4 (Figure 4F). Finally, through the evaluation results of all normalization methods, we find that although the Z-score method is relatively ideal for QC data correction, it shows great instability among samples as shown in Figure 4H.

**Figure 4.**
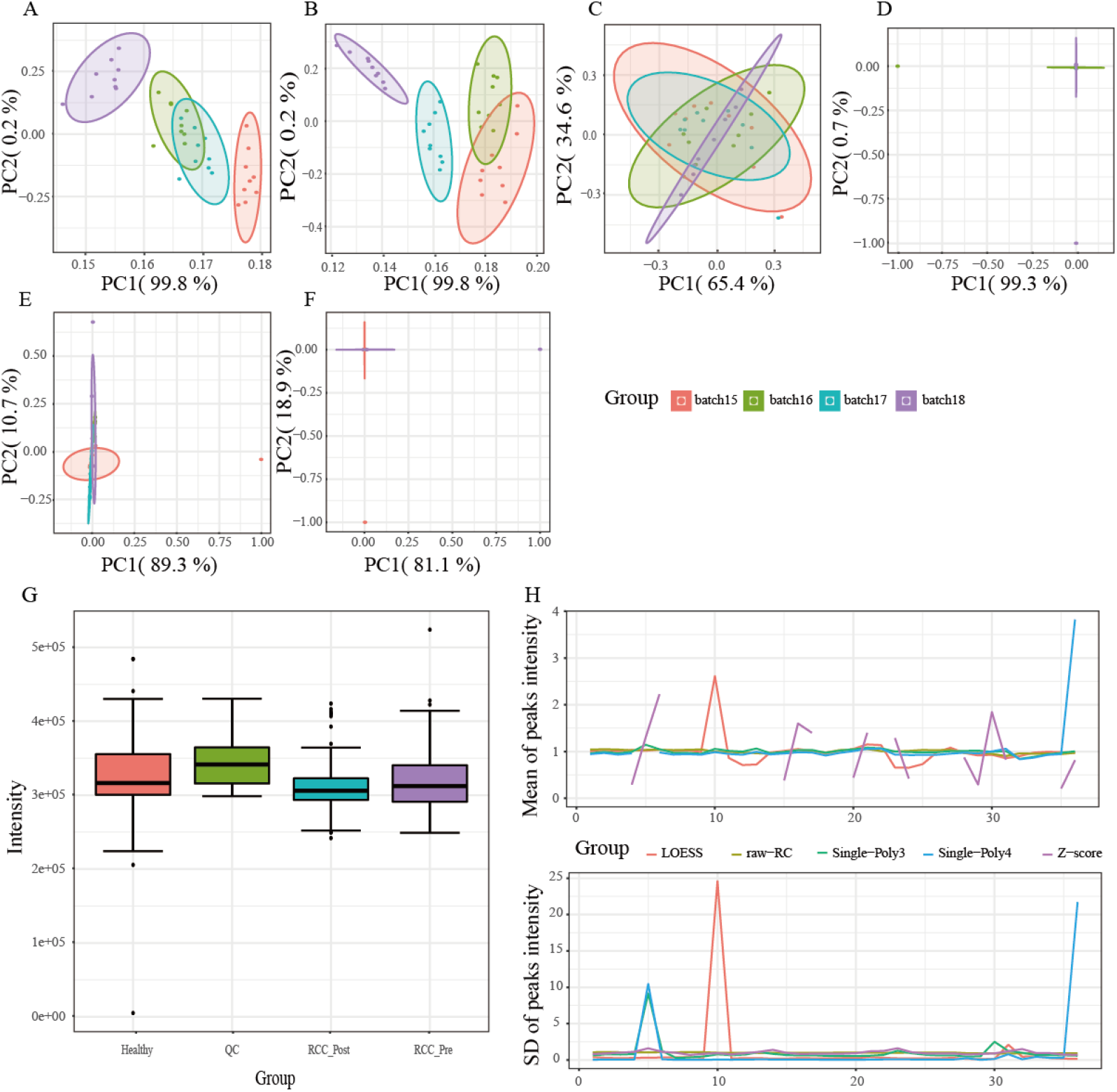
Normalization for QC data by Normalization pipeline 1. A-F present the PCA results for QC data. A) Raw QC data. B) Mass intensity data adjusted by RC. C) Mass intensity data was adjusted by QC based on Z-score. D) Mass intensity data adjusted by QC based on LOESS. E) Mass intensity data adjusted by QC based on Single-Poly3. F) Mass intensity data adjusted by QC based on Single-Poly4. G) RC peak intensity distribution by groups. H) Comparison of all normalization methods.

**Figure 5.**
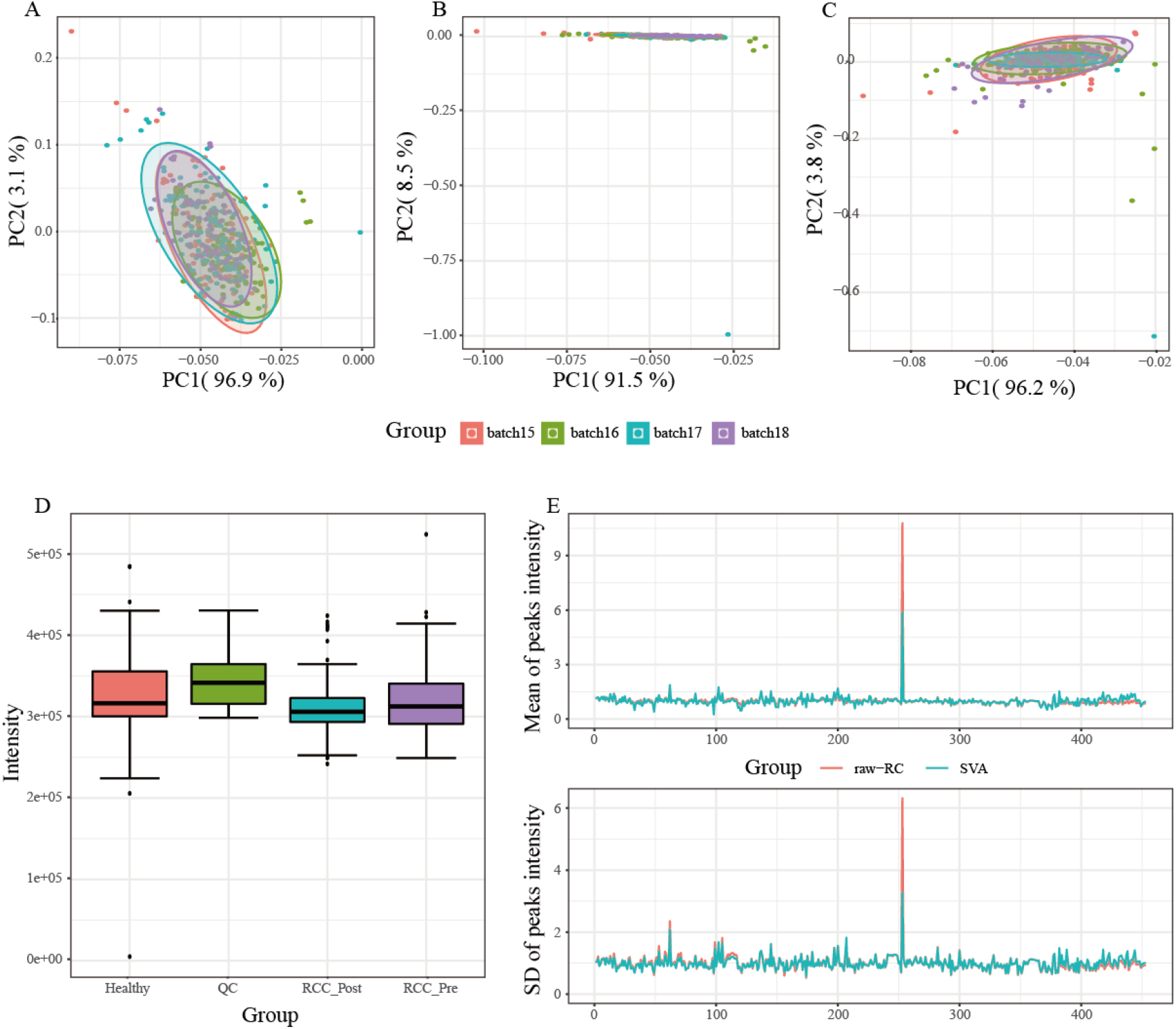
Normalization for RCC data using normalization pipeline 2. A-C present the PCA results for sample intensity data. A) Raw sample data. B) Sample data adjusted by RC. C) Sample data mass intensity data adjusted by SVA. D) RC peak intensity distribution by groups. E) Comparison of all normalization methods.

To verify the accuracy of our normalization pipeline 1, we use normalization pipeline 2 to explore whether the data can be normalized by the machine learning algorithm. From Figure 5, we can see that the raw data did not have any clustering distribution between groups (Figure 5A), and the results based on the RC (Figure 5B) and SVA algorithms (Figure 5C) still did not adjust the difference distribution between groups. Meanwhile, we checked the difference between raw-RC data and the RC data adjusted by the machine learning algorithm and found that they were no different from each other (Figure 5E). Therefore, we believe that there is an experimental error in the raw experimental data, and the subsequent relevant analysis cannot be carried out.

Of course, the above results were based on the experimental data that we thought had problems. Next, we repeated the normalization pipeline 2 using the previously published data(Li, et al., 2019) of the laboratory to further prove the problems of the RCC data. Here, positive ion data from 120 patients with colorectal cancer and 120 healthy controls were introduced for analysis. As this part of the data is only corrected based on RC, only the analysis results of normalization pipeline 2 were presented. From the results, we found that there was no difference in RC between the two groups of data (Figure 6C). Meanwhile, this part of the data is obviously clustered into two groups of samples, which is the same as our raw grouping (Figure 6A). To eliminate the influence of the batch, we carried out normalization based on RC and SVA and found that it was consistent with the raw data, indicating that batch also had little influence on the data (Figure 6B and C). Finally, after comparing the raw data with the data processed by SVA, it was found that the raw data were consistent with the normalized data (Figure 6D).

**Figure 6.**
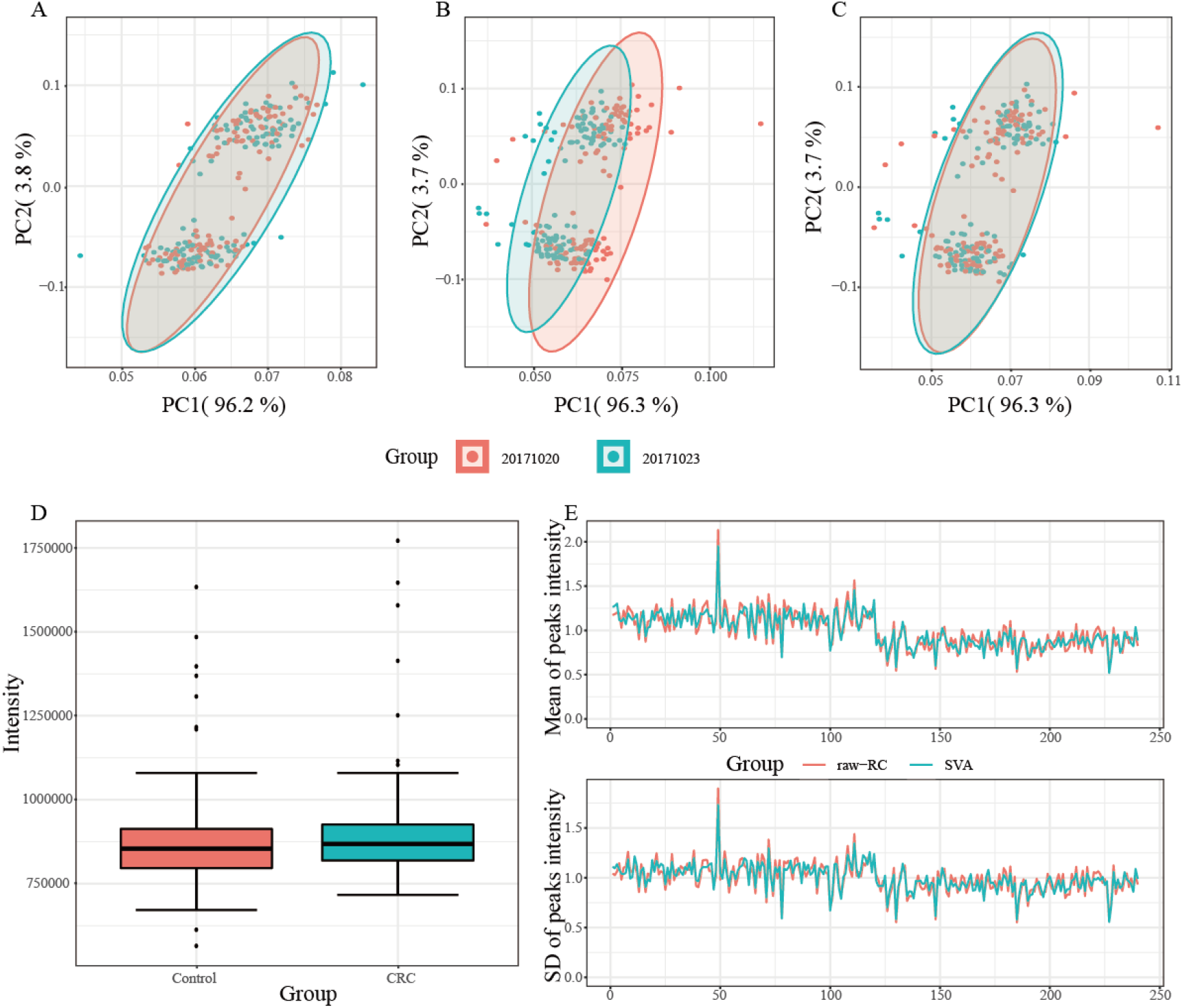
Normalization for CRC data by using normalization pipeline 2. Refer to Figure 5 for legend details.

Finally, to verify the consistency between our MARC analysis process and previously published results, we run analysis process 2 to conduct an in-depth analysis of CRC data, as shown in Figure 7. In this analysis process, we analyzed the differential metabolites between CRC and healthy samples using the Wilcoxon test (Figure 7A) and OPLS-DA (Figure 7B) methods and found 98 peaks in both analyses (Figure 7C). To verify the reliability of our refMet database, we screened all databases to explore the metabolite distribution for 98 differential ion peaks and found that more metabolites were matched in the MASSBANK and HMDB databases (Figure 7D).

**Figure 7.**
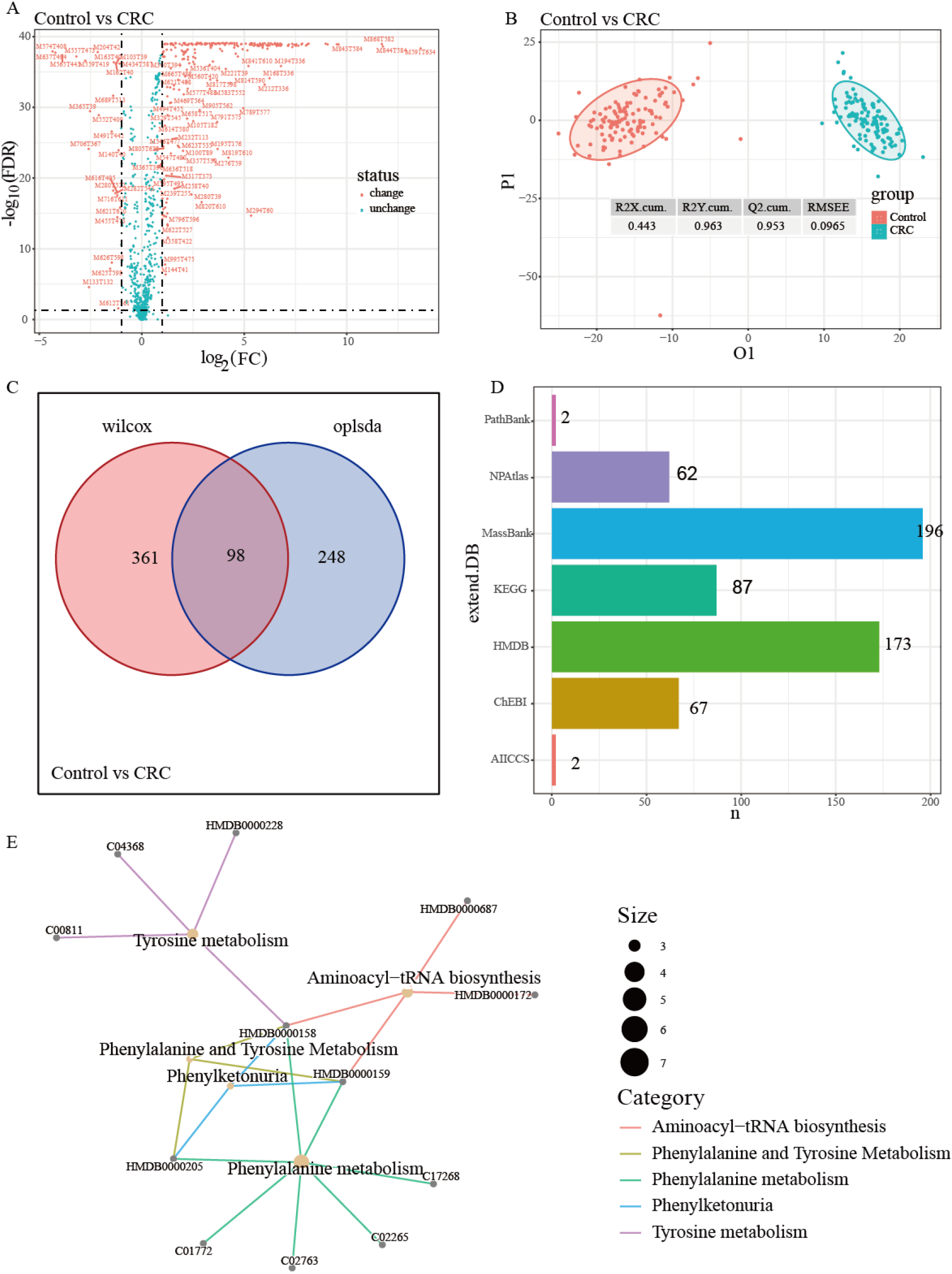
Analysis process pipeline 2 results. A) Differential analysis using the Wilcox test. B) Differential analysis using the OPLS-DA. C) Differential metabolic ion peak number statistics of A and B results. D) Distribution of 98 differential metabolic ion peaks in all databases of our refMet. E) Pathway enrichment analysis by 98 metabolic peaks.

Finally, we summarized the existing related software and compared the software with 8 advantages of our software, including data upload, visual operation, cross-platform, localized configuration, analysis report construction, automatic process analysis, complete analysis strategy, and refMet database (Table 1). Due to the limitation of the web browser, the upload of large files exceeds the wait time of the network and thus the uploading is interrupted, which has become the limitation of many platforms on the sample size. In the existing analysis software, W4M has no restriction on data uploading due to the analysis platform built on the Galaxy platform (https://galaxyproject.org/). Visual operation facilitates more users to achieve metabolic data analysis with the mouse. Numerous softwares deployed in the cloud can achieve visual operation but not for mzMatch/PeakML. Cross-platform means that software is not limited by the operating system and hardware environment. Cloud-based software generally only supports a developer’s remote server environment, of course, if the source code is open access, it can also be deployed locally to achieve cross-platform use such as W4M. Local configuration refers to whether the software can be run on a user’s machine. Analysis report construction refers to whether the software can achieve a standardized report summary of data results. Automatic process analysis refers to whether the software can simplify operations and automate analysis, which is a beginner-friendly procedure without requiring complicated parameter settings. The complete analysis strategy refers to whether the analysis of untargeted metabolic data is a complete analysis process from the raw data to the differential metabolic ions, and there are corresponding analysis strategies for each analysis node. refMet refers to the common data set used in the identification of metabolic ions. The reliability of refMet plays an important role in later metabolite verification. From Table 1, we found that only MetaboAnalyst and openNAU had high functional consistency.

**Table 1.** Comparison of metabolic data analysis software. **✗** represent software does not have this function, **✓** is used as the opposite

| Function | MetaboAnalyst | mzMatch/<br>PeakML | GNPS | NOREVA | W4M | Metabolomics<br>Workbench |
| --- | --- | --- | --- | --- | --- | --- |
| Upload<br>limitation | ✕ | ✕ | ✕ | ✕ | ✓ | ✕ |
| Visual<br>operation | ✓ | ✓ | ✓ | ✓ | ✓ | ✓ |
| Cross-platform | ✓ | ✓ | ✗ | ✗ | ✗ | ✗ |
| Localized configuration | ✓ | ✓ | ✗ | ✗ | ✓ | ✗ |
| Analysis report construction | ✓ | ✗ | ✓ | ✓ | ✗ | ✗ |
| Automatic process analysis | ✗ | ✗ | ✗ | ✗ | ✗ | ✗ |
| Complete analysis strategy | ✓ | ✗ | ✗ | ✗ | ✓ | ✗ |
| refMet database | ✓ | ✗ | ✗ | ✓ | ✓ | ✓ |

To compare the differences between the two softwares, we uploaded the raw peak intensity data and normalized peak data based on SVA of the CRC data to MetaboAnalyst 5.0 and performed functional analysis. First, we used the data analysis strategy (Normalization by reference feature and Auto scaling) of MetaboAnalyst 5.0 to normalize CRC raw ion peak data, and the results before and after normalization. Then, we obtained the results (5 pathways and 104 peaks) of enrichment analysis (mummichog and GSEA, Supplement Table S1a) and differential metabolites based on normalized ion peak data. To compare the differences between our analysis strategies and MetaboAnalyst 5.0, we submitted our SVA normalized peak data and did not perform any normalization, which showed 3 enriched pathways (Supplement Table S1b) and 141 differential ion peaks. In openNAU, we found 98 different metabolic ion peaks, which were enriched in 5 related pathways (Figure 7E). By comparing the above results, we found that the SVA data based on our analysis strategy cannot obtain meaningful results in MetaboAnalyst 5.0. In addition, we performed pathway enrichment analysis based on 98 metabolic ions (positive ions, Figure 7E) and 61 differential metabolites (positive ions and negative ions, refers to Supplementary Figure 1C(Li, et al., 2019)) identified by secondary mass spectrometry between CRC and healthy samples and found a high consistency for pathways. But only 3 consistent pathway results in MetaboAnalyst 5.0 pathway enrichment analysis. This also shows the reliability of our RefMet database and software.

## Discussion

We constructed an open-source complete analysis platform of untargeted metabolomics from raw data to differential metabolites. Because of the privacy and security of raw data in metabolomics research, we divided the software into two parts: localization pretreatment and cloud data analysis. Certainly, the localized part can also be directly deployed in the cloud to achieve a seamless connection with the cloud data analysis. In addition, we built an API of the MetaboAnalyst software, which can directly convert our preprocessing results into its uploaded data format (https://www.metaboanalyst.ca/MetaboAnalyst/upload/PeakUploadView.xhtml). In our software, we start with an analysis strategy, focus on analysis process pipelines, integrate current untargeted metabolomics-related software and other comics-related method tools, and build analysis strategy and analysis process for users. From the extraction of raw data, quality control, and correction to every analysis process, we provide a complete analysis strategy to transform the complex analysis process into simple visual human-machine communication.

Z score as a normalization method for decentralization, describes a value’s relationship to the mean and standard deviation of a group of values. In our software, we normalized each ion of the experimental sample to obtain the Z score based on the mean and standard deviation of the quality control group. We found that raw peak data can be well corrected even in experimental data of poor quality (Figure 4C). Of course, this method also has certain limitations, that is, without quality control samples, it cannot be scored. We think this approach may be more general in normalizing untargeted metabolic data based on quality control samples.

Obviously, MS-DIAL software(Tsugawa, et al., 2015) has has developed a complete metabolomic ion extraction and data analysis process. However, in the data normalization module, MS-DIAL only performed LOESS and internal standard ions elimination methods. Statistical analysis of MS-DIAL was performed based on EXCEL software macro (http://prime.psc.riken.jp/compms/others/main.html), including Principal Component Analysis (PCA), projection to Latent Structure regression (PLS-R), and projection to latent structure based discriminant analysis (PLS-DA) analysis. We found that the normalized approach was too simple to cope with the growing number of complex metabolomics data, and the cost of Microsoft Office was not negligible. Meanwhile, the operating efficiency and operating environment of EXCEL software are limited by computer configuration and system. openNAU was based on the open source R and PHP languages for the visual operation of untargeted metabolic data analysis, providing more space and cost savings for users to choose software. In addition, our enrichment analysis strategy is superior to MetaboAnalyst5.0 in untargeted metabolomics analysis.

We hope to build analysis processes for different scenarios and summarize and annotate them in text form. Based on the label of each pipeline, we can build artificial intelligence to create complex analysis processes based on simple human-machine communication(Li, et al., 2021).

Of course, the platform still has some shortcomings in the analysis process. From the perspective of calculation time, since the IPO software will spend considerable time in the process of optimizing parameters, the time spent in the process of preprocessing raw data will increase exponentially with increasing of data volume. Compared with MS-DIAL software, we need to continue to improve the MetaQC module to expand the data types of untargeted metabolomics, refine ion extraction parameters, and increase individual operations for imported samples. In addition, we found Norm ISWSVR as a recently released package favorably improves the data quality of large-scale metabolomics data(Ding, et al., 2022). We will overcome this shortcoming and add new analysis packages in the future.

## Conclusions

In this article, we constructed the OpenNAU software for untargeted metabolic data evaluation and analysis platforms. The software will be the first easy-to-use analytics platform with analytics processes at its core. In the meantime, the software effectively protects raw data while lowering the difficulty for untargeted metabolomics analysis. In addition, it not only provides the data quality assessment process but also provides the complete process of untargeted metabolomics analysis.

## Acknowledgments

The authors appreciate the scholarly comments of Dr. Jing Li, China Pharmaceutical University. This work was supported by Alibaba Cloud.

## Contributions

Q.S. developed the protocol for OpenNAU, with help from M.W. M.L., D.Z., Y.W., Q.S., Q.X. and M.W. wrote and revised the manuscript. M.L and Q.S conceived the original idea and designed the study. Q.X contributed to the experiments. All authors agreed with the conclusion and approved the final version of manuscript.

## Supplementary data

**Supplement Table S1.** Functional analysis results of MetaboAnalyst 5.0 based on raw peak intensity data and normalized data using SVA in openNAU. The threshold value was Combined_Pvals<0.05.

| a) Pathway results for raw CRC data |  |  |  |
| --- | --- | --- | --- |
| Pathway | Mummichog_Pvals | GSEA_Pvals | Combined_Pvals |
| Tyrosine metabolism | 0.1415 | 0.01075 | 0.01139 |
| Tryptophan metabolism | 0.276 | 0.01351 | 0.02458 |
| Glycerophospholipid metabolism | 0.5534 | 0.01031 | 0.03518 |
| De novo fatty acid biosynthesis | 0.54 | 0.01075 | 0.03569 |
| Arachidonic acid metabolism | 0.276 | 0.02703 | 0.044 |

| b) Pathway results for CRC data normalized by SVA |  |  |  |
| --- | --- | --- | --- |
| Pathway | Mummichog_Pvals | GSEA_Pvals | Combined_Pvals |
| Linoleate metabolism | 0.1555 | 0.01163 | 0.01323 |
| Biopterin metabolism | 0.216 | 0.01471 | 0.02145 |
| Tyrosine metabolism | 0.7693 | 0.01064 | 0.04752 |

